# Phenome-wide association analysis of LDL-cholesterol lowering genetic variants in *PCSK9*

**DOI:** 10.1101/329052

**Authors:** Amand F Schmidt, Michael V Holmes, David Preiss, Daniel Swerdlow, Spiros Denaxas, Ghazaleh Fatemifar, Rupert Faraway, Chris Finan, Tom Lumbers, Albert Henry, Dennis Valentine, Zammy Fairhurst-Hunter, Fernando Pires Hartwig, Bernardo Lessa Horta, Elina Hypponen, Christine Power, Max Moldovan, Erik van Iperen, Kees Hovingh, Ilja Demuth, Kristina Norman, Elisabeth Steinhagen-Thiessen, Juri Demuth, Lars Bertram, Christina M Lill, Stefan Coassin, Johann Willeit, Stefan Kiechl, Karin Willeit, Dan Mason, John Wright, Richard Morris, Goya Wanamethee, Peter Whincup, Yoav Ben-Shlomo, Stela McLachlan, Jackie F. Price, Mika Kivimaki, Catherine Welch, Adelaida Sanchez-Galvez, Pedro Marques-Vidal, Andrew Nicolaides, Andrie G. Panayiotou, N. Charlotte Onland-Moret, Yvonne T. van der Schouw, Giuseppe Matullo, Giovanni Fiorito, Simonetta Guarrera, Carlotta Sacerdote, Nicholas J Wareham, Claudia Langenberg, Robert A Scott, Jian’an Luan, Martin Bobak, Sofia Malyutina, Andrzej Pajak, Ruzena Kubinova, Abdonas Tamosiunas, Hynek Pikhart, Niels Grarup, Oluf Pedersen, Torben Hansen, Allan Linneberg, Tine Jess, Jackie Cooper, Steve E Humphries, Murray Brilliant, Terrie Kitchner, Hakon Hakonarson, David S. Carrell, Catherine A. McCarty, Kirchner H Lester, Eric B. Larson, David R. Crosslin, Mariza de Andrade, Dan M Roden, Joshua C Denny, Cara Carty, Stephen Hancock, John Attia, Elizabeth Holliday, Rodney Scott, Peter Schofield, Martin O’Donnell, Salim Yusuf, Michael Chong, Guillaume Pare, Pim van der Harst, M. Abdullah Said, Ruben N. Eppinga, Niek Verweij, Harold Snieder, Tim Christen, D.O. Mook-Kanamori, Stefan Gustafsson, Lars Lind, Erik Ingelsson, Raha Pazoki, Oscar Franco, Albert Hofman, Andre Uitterlinden, Abbas Dehghan, Alexander Teumer, Sebastian Baumeister, Marcus Dörr, Markus M. Lerch, Uwe Völker, Henry Völzke, Joey Ward, Jill P Pell, Tom Meade, Ingrid E. Christophersen, Anke H. Maitland-van der Zee, Ekaterina V. Baranova, Robin Young, Ian Ford, Archie Campbell, Sandosh Padmanabhan, Michiel L Bots, Diederick E. Grobbee, Philippe Froguel, Dorothée Thuillier, Ronan Roussel, Amelie Bonnefond, Bertrand Cariou, Melissa Smart, Yanchun Bao, Meena Kumari, Anubha Mahajan, Jemma C. Hopewell, Sudha Seshadri, Caroline Dale, Rui Providencia E Costa, Paul M Ridker, Daniel I. Chasman, Alex P. Reiner, Marylyn D Ritchie, Leslie A Lange, Alex J. Cornish, Sara E. Dobbins, Kari Hemminki, Ben Kinnersley, Marc Sanson, Karim Labreche, Matthias Simon, Melissa Bondy, Philip Law, Helen Speedy, James Allan, Ni Li, Molly Went, Niels Weinhold, Gareth Morgan, Pieter Sonneveld, Björn Nilsson, Hartmut Goldschmidt, Kari Hemminki, Amit Sud, Andreas Engert, Markus Hansson, Harry Hemingway, Folkert W Asselbergs, Riyaz S Patel, Brendan J Keating, Naveed Sattar, Richard Houlston, Juan P Casas, Aroon D Hingorani

## Abstract

**Background:** We characterised the phenotypic consequence of genetic variation at the *PCSK9* locus and compared findings with recent trials of pharmacological inhibitors of PCSK9.

**Methods:** Published and individual participant level data (300,000+ participants) were combined to construct a weighted *PCSK9* gene-centric score (GS). Fourteen randomized placebo controlled PCSK9 inhibitor trials were included, providing data on 79,578 participants. Results were scaled to a one mmol/L lower LDL-C concentration

**Results:** The *PCSK9* GS (comprising 4 SNPs) associations with plasma lipid and apolipoprotein levels were consistent in direction with treatment effects. The GS odds ratio (OR) for myocardial infarction (MI) was 0.53 (95%CI 0.42; 0.68), compared to a PCSK9 inhibitor effect of 0.90 (95%CI 0.86; 0.93). For ischemic stroke ORs were 0.84 (95%CI 0.57; 1.22) for the GS, compared to 0.85 (95%CI 0.78; 0.93) in the drug trials. ORs with type 2 diabetes mellitus (T2DM) were 1.29 (95% CI 1.11; 1.50) for the GS, as compared to 1.00 (95%CI 0.96; 1.04) for incident T2DM in PCSK9 inhibitor trials. No genetic associations were observed for cancer, heart failure, atrial fibrillation, chronic obstructive pulmonary disease, or Alzheimer’s disease – outcomes for which large-scale trial data were unavailable.

**Conclusions:** Genetic variation at the *PCSK9* locus recapitulates the effects of therapeutic inhibition of PCSK9 on major blood lipid fractions and MI. Apparent discordance between genetic associations and trial outcome for T2DM might be explained lack by a of statistical precision, or differences in the nature and duration of genetic versus pharmacological perturbation of PCSK9.

**Funding:** This research was funded by the British Heart Foundation (SP/13/6/30554, RG/10/12/28456, FS/18/23/33512), UCL Hospitals NIHR Biomedical Research Centre, by the Rosetrees and Stoneygate Trusts.

**Condensed abstract:** Evidence on the long-term efficacy and safety of therapeutic inhibition of PCSK9 is lacking. To explore potential long-term effects of PCSK9 inhibition, we characterised the phenotypic consequence of LDL-cholesterol lowering variants at the *PCSK9* locus. A *PCSK9* gene score comprising 4 SNPs recapitulated the effects of therapeutic inhibition of PCSK9 on major blood lipid fractions and risk of myocardial infarction, and was associated with an increased risk of type 2 diabetes. No associations with safety outcomes such as cancer, COPD, Alzheimer’s disease or atrial fibrillation were identified. Our findings suggest PCSK9 inhibition may be safe and effective during prolonged use.

## Introduction

Statins and ezetimibe reduce the risk of major coronary events and ischemic stroke via lowering of low density lipoprotein-cholesterol (LDL-C) (1–3). Loss-of-function mutations in *PCSK9* are associated with lower LDL-C and a reduced risk of coronary heart disease (CHD)(4, 5). Antibodies (mAbs) inhibiting PCSK9, reduce LDL-C in patients with hypercholesterolaemia, and received market access in 2015. The FOURIER and ODYSSEY OUTCOMES trials tested the efficacy of PCSK9-inhibition versus placebo on the background of statin treatment and both found that PCSK9 inhibition led to a 15% relative risk reduction of major vascular events in patients with established CVD and recent acute coronary syndrome over a median follow up of 2.2 or 2.8 years(6, 7).

Evidence is limited on the effect of PCSK9 inhibition on clinical outcomes, and on safety outcomes that might only become apparent with prolonged use. Nor is evidence available on the efficacy and safety of PCSK9 inhibitors in subjects other than the high-risk patients studied in trials. Mendelian randomisation for target validation uses naturally-occurring variation in a gene encoding a drug target to identify mechanism-based consequences of pharmacological modification of the same target(8). Such studies have previously proved useful in predicting success and failure in clinical trials and have assisted in delineating on-target from off-target actions of first-in-class drugs(9–13). For example, previous studies showed that variants in *HMGCR,* encoding the target for statins, were associated with lowering concentrations of LDL-C and lower risk of coronary heart disease(9) (CHD), while confirming the on-target nature of the effect of statins on higher body weight and higher risk of type 2 diabetes (T2DM)(9).

We characterised the phenotypic consequences of genetic variation at *PCSK9* in a large, general population sample focussing on therapeutically relevant biomarkers, cardiovascular disease (CVD), individual CVD components and non-CVD outcomes such as cancer, Alzheimer’s disease, and chronic obstructive pulmonary disease (COPD). Effect estimates from the genetic analysis were compared to those from intervention trials where the outcomes under evaluation overlapped.

## Methods

We summarise methods briefly here as they have been previously described in detail(14).

### Genetic variant selection

SNPs rs11583680 (minor allele frequency [MAF] = 0.14), rs11591147 (MAF = 0.01), rs2479409 (MAF = 0.36) and rs11206510 (MAF = 0.17) were selected as genetic instruments at the *PCSK9* locus based on the following criteria: (1) an LDL-C association as reported by the Global Lipids Genetics Consortium (GLGC)^14^; (2) low pairwise linkage disequilibrium (LD) (r^2^ ≤0.30) with other SNPs in the region (based on 1,000 Genomes CEU data); and (3) the combined annotation dependent depletion (CADD) score(15) which assesses potential functionality (see Appendix table 1).

Previously, we explored the between-SNP correlations (see Appendix figure 1 of Schmidt et al 2017(14)), revealing an r-squared of 0.26 between rs11206510 and rs11583680, confirming all other SNPs were approximately independent (r^2^ ≤ 0.07). Subsequent adjustment for the residual LD (correlation) structure did not impact results (see Appendix figure 90 of Schmidt et al 2017(14)).

### Individual participant-level and summary-level data

Participating studies (Appendix table 2) provided analyses of individual participant-level data (IPD) based on a common analysis script (available from AFS), submitting summary estimates to the UCL analysis centre. These data were supplemented with public domain data from relevant genetic consortia (Appendix table 3). Studies contributing summary estimates to genetic consortia were excluded from the IPD component of the analysis to avoid duplication.

Biomarker data were collected on the major routinely measured blood lipids (LDL-C, HDL-C, triglycerides [TG], total cholesterol [TC]); apolipoproteins A1 [ApoA1] and B [ApoB], and nominal lipoprotein (Lp)(a); systolic (SBP) and diastolic (DBP) blood pressure; inflammation markers C-reactive protein (CRP), interleukin-6 (IL-6), and fibrinogen; haemoglobin; glycated haemoglobin (HbA_1c_); liver enzymes gamma-glutamyltransferase (GGT), alanine aminotransferase (ALT), aspartate transaminase (AST), and alkaline phosphatase (ALP); serum creatinine, and cognitive function (standardized to mean 0, and standard deviation 1, see Appendix table 5).

We focussed on individual clinical endpoints, rather than composites, which have been assessed in outcome trials, as well as disease end-points commonly seen in patients likely to be eligible for PCSK9 inhibitor treatment. Ischemic CVD endpoints studied were myocardial infarction (MI), ischemic stroke, revascularization, and angina. The following non-ischemic CVD events were considered: haemorrhagic stroke, heart failure, and atrial fibrillation. Non-CVD outcome data was collected on common chronic diseases: COPD, any cancer (including those of the breast, prostate, colon and lung), Alzheimer’s disease, and T2DM.

Finally, aggregated trial data on the effect of monoclonal PCSK9 inhibitors were compared to placebo for MI, revascularization, ischemic or haemorrhagic stroke, cancer, and T2DM abstracted from the Cochrane systematic review(6). We compared effects on biomarkers and clinical endpoints common to both the genetic analysis and trials.

### Statistical analyses

In all analyses, we assumed an additive allele effect with genotypes coded as 0, 1 and 2, corresponding to the number of LDL-C lowering alleles. Continuous biomarkers were analysed using linear regression and binary endpoints using logistic regression. Study-specific associations were pooled for each SNP using the inverse variance weighted method for fixed effect meta-analysis. Study-specific associations were excluded if the SNP was not in Hardy-Weinberg equilibrium (see Appendix Table 4, based on a Holm-Bonferroni alpha criterion). We estimated the effect at the *PCSK9* locus by combining all four SNPs in a gene centric score (GS).

Trial data were assembled as per Schmidt et al. 2017(6). Briefly, systematic searches were performed using the Cochrane Central Register of Controlled Trials (CENTRAL), MEDLINE, Embase, Web of Science registries, Clinicaltrials.gov and the International Clinical Trials Registry Platform databases. Data from placebo controlled trials were extracted and combined using the inverse variance weighted method for continuous data and a random-intercept logistic regression model for binary data(6).

Results are presented as mean differences (MD) or odds ratios (OR) with 95% confidence intervals (CI). Analyses were conducted using the statistical programme R version 3.4.1(16).

### Role of the funding source

The funder(s) of the study played no role in study design, data collection, data analysis, data interpretation, or writing of the report. AFS had full access to all the data in the study and shares final responsibility for the decision to submit for publication with all authors.

Local ethics committees for studies contributing data to these analyses granted approval for the work.

## Results

Participant level data were available from up to 246,355 individuals, and were supplemented by summary effect estimates from data repositories, resulting in a sample size of 320,170 individuals, including 95,865 cases of MI, 16,437 stroke, 11,920 ischemic stroke, 51,623 T2DM, 54,702 cancer, 25,630 Alzheimer’s disease and 12,412 of COPD.

### Lipid and apolipoprotein associations

As reported previously(14), the four *PCSK9* SNPs were associated with lower LDL-C blood concentrations ranging from −0.02 mmol/L (95%CI −0.03; −0.02) per allele for rs11583680 to −0.34 mmol/L (95%CI −0.36; −0.32) for rs11591147 (See Appendix Figure 1). *PCSK9* SNPs associated with a lower LDL-C concentration were also associated with lower concentrations of apolipoprotein B proportionate to the LDL-C association.

Associations of the GS with the other lipids or apolipoproteins, scaled to a 1 mmol/L lower LDL-C were (Table 1): 0.05 mmol/L (95%CI 0.02; 0.07) for HDL-C, −0.07 mmol/L (95%CI −0.12; −0.01) for TG, −1.06 mmol/L (95%CI −1.12; −1.00) for TC, −0.20 g/L (95%CI −0.25; -0.18) for ApoB, 0.02 g/L (95%CI −0.01; 0.06) for ApoA1, and −4.12 mg/dL (95%CI −8.62; 0.38) for Lp(a).

**Table 1.**
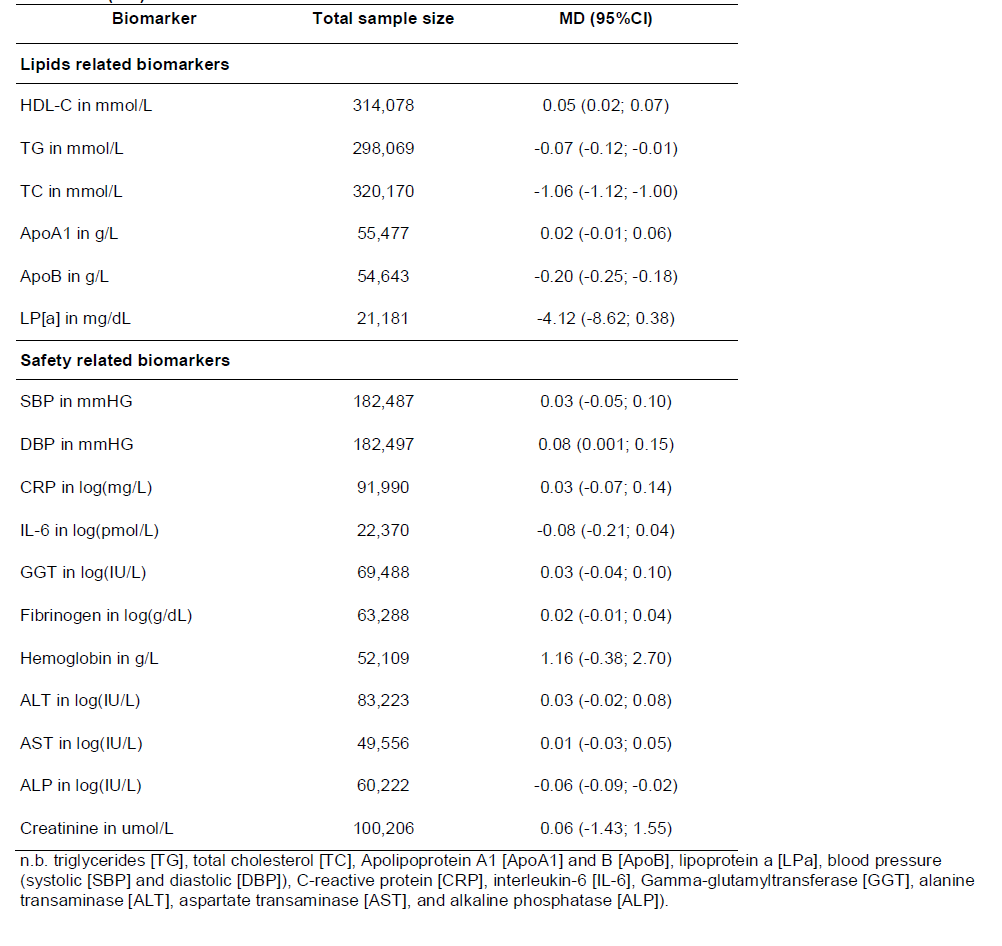
Biomarker associations of a *PCSK9* gene centric score, effect presented as mean difference (MD) with 95% confidence interval in brackets with the effects scaled to a 1 mmol/L decrease in LDL-C.

| Biomarker | Total sample size | MD (95%CI) |
| --- | --- | --- |
| **Lipids related biomarkers** |  |  |
| HDL-C in mmol/L | 314,078 | 0.05 (0.02; 0.07) |
| TG in mmol/L | 298,069 | -0.07 (-0.12; -0.01) |
| TC in mmol/L | 320,170 | -1.06 (-1.12; -1.00) |
| ApoA1 in g/L | 55,477 | 0.02 (-0.01; 0.06) |
| ApoB in g/L | 54,643 | -0.20 (-0.25; -0.18) |
| LP[a] in mg/dL | 21,181 | -4.12 (-8.62; 0.38) |
| **Safety related biomarkers** |  |  |
| SBP in mmHG | 182,487 | 0.03 (-0.05; 0.10) |
| DBP in mmHG | 182,497 | 0.08 (0.001; 0.15) |
| CRP in log(mg/L) | 91,990 | 0.03 (-0.07; 0.14) |
| IL-6 in log(pmol/L) | 22,370 | -0.08 (-0.21; 0.04) |
| GGT in log(IU/L) | 69,488 | 0.03 (-0.04; 0.10) |
| Fibrinogen in log(g/dL) | 63,288 | 0.02 (-0.01; 0.04) |
| Hemoglobin in g/L | 52,109 | 1.16 (-0.38; 2.70) |
| ALT in log(IU/L) | 83,223 | 0.03 (-0.02; 0.08) |
| AST in log(IU/L) | 49,556 | 0.01 (-0.03; 0.05) |
| ALP in log(IU/L) | 60,222 | -0.06 (-0.09; -0.02) |
| Creatinine in umol/L | 100,206 | 0.06 (-1.43; 1.55) |

n.b. triglycerides [TG], total cholesterol [TC], Apolipoprotein A1 [ApoA1] and B [ApoB], lipoprotein a [LPa], blood pressure (systolic [SBP] and diastolic [DBP]), C-reactive protein [CRP], interleukin-6 [IL-6], Gamma-glutamyltransferase [GGT], alanine transaminase [ALT], aspartate transaminase [AST], and alkaline phosphatase [ALP]).

The associations of the *PCSK9* GS with blood-based lipid markers were directionally concordant with effects from treatment trials of therapeutic inhibition of PCSK9 (Figure 1).

**Figure 1.**
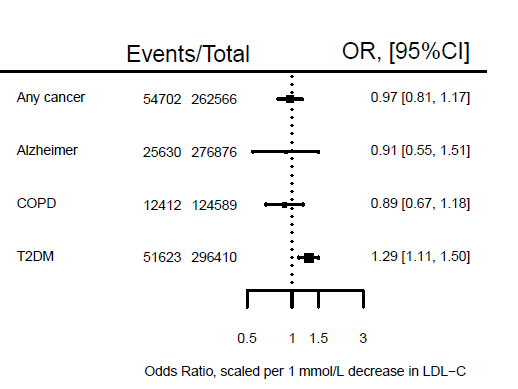
Lipid and lipoprotein associations of a *PCSK9* gene-centric score (GS) compared to placebo-controlled randomized trials of therapeutic inhibition of PCSK9 Footnote: Effect estimates are presented as mean differences, with 95% confidence interval (CI). Trial estimates are presented as percentage change from baseline, and GS estimates scaled to a 1 mmol/L lower LDL-C (mmol/L). Results are pooled using a fixed effect model. Trial estimates are based on the systematic review by Schmidt *et al* 2017(6)

### Genetic associations with other biochemical and physiological measures

The GS estimates with SBP and DBP were 0.03 mmHg (95%CI −0.05; 0.10) and 0.08 mmHg (95%CI 0.0001; 0.15), respectively, per 1 mmol/L lower LDL-C. The *PCSK9* GS was associated with nominally lower ALP (IU/L) −0.06 (95%CI −0.09; −0.02), but not with other liver enzymes (Table 1).

### Genetic associations with ischemic cardiovascular events

The *PCSK9* GS was associated with a lower risk of MI (OR 0.53; 95%CI 0.42; 0.68; 95,865 cases), which was directionally consistent with results from placebo-controlled PCSK9 inhibition trials: OR 0.90 (95%CI 0.86; 0.93), with both estimates scaled to a 1 mmol/L lower LDL-C (Figures 2 and 3). The genetic effect estimate for ischemic stroke was OR 0.84 (95%CI 0.57; 1.22; 11,920 cases), concordant in direction to that of the drugs trials (OR 0.85 95%CI 0.78; 0.93). Similarly, the *PCSK9* GS association with coronary revascularization (OR 0.75 95%CI 0.44; 1.27) was directionally consistent with the PCSK9 inhibitor trials (OR 0.90; 95%CI 0.86, 0.93) (Figure 3).

**Figure 2.**
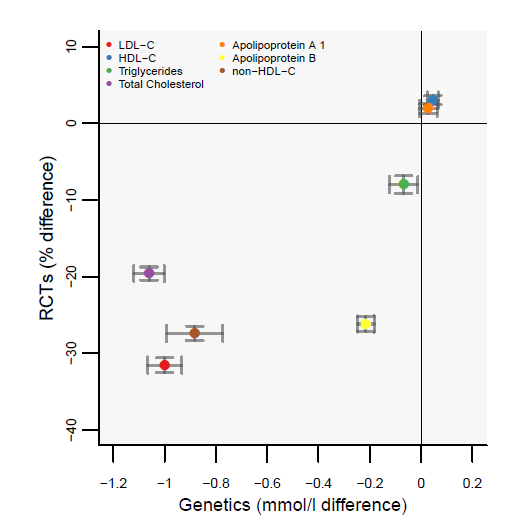
Associations of a *PCSK9* gene-centric score with ischemic and non-ischemic cardiovascular endpoints. Footnote: Effect estimates are presented as odds ratios (OR), with 95% confidence interval (CI) scaled to a 1 mmol/L lower LDL-C (mmol/L). Results are pooled using a fixed effect model. The size of the squares are proportional to the inverse of the variance.

**Figure 3.**
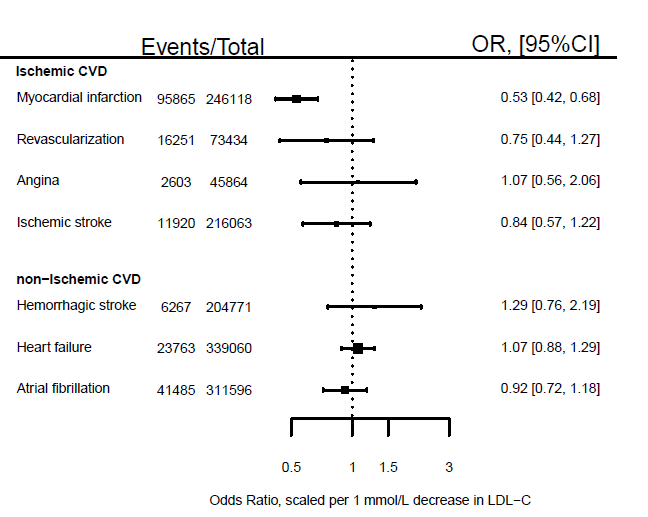
Clinical endpoint associations of the *PCSK9* gene-centric score (GS) as compared to placebo-controlled randomized trials of therapeutic inhibition of PCSK9. Footnote: Effect estimates are presented as odds ratios (OR), with 95% confidence interval (CI), for the GS scaled to a 1 mmol/L lower LDL-C (mmol/L). Results are pooled using a fixed effect model. Trial estimates are based on the systematic review by Schmidt *et al* 2017(6), with the estimates on ischemic stroke and revascularization solely based on the FOURIER and ODYSSEY OUTCOMES trials.

### Genetic associations with non-ischemic cardiovascular disease

The point estimate for the GS association with hemorrhagic stroke (Figure 2), OR 1.29 (95%CI 0.76; 2.19), was discordant to the estimate from PCSK9 inhibitor trials (OR 0.96 95%CI 0.75; 1.23) (Figure 3), although the confidence intervals overlapped. Comparing the association of *PCSK9* GS with hemorrhagic and ischemic stroke indicated the GS had a differential effect (p-value = 0.02). No *PCSK9* GS association was observed with atrial fibrillation (OR 0.92 95%CI 0.72; 1.18; 41 485 cases), or heart failure (OR 1.06 95%CI 0.88; 1.28; 23,763 cases) (Figure 2).

### Associations with non-cardiovascular disease and related biomarkers

The *PCSK9* GS was not associated with the risk of any cancer (OR 0.97: 95%CI 0.81; 1.17; 54,702 cases, see Figure 4), nor with any of 13 specific types of cancer (Appendix Figure 2). We did not observe an association with either Alzheimer’s disease or cognitive performance: for Alzheimer’s the OR was 0.91 (95%CI 0.55; 1.51) and for cognition (per standard deviation) −0.03 (95%CI −0.22; 0.16). As reported before(14) the GS was associated with T2DM (OR 1.00 95%CI 1.11; 1.50) (Figure 4), higher body weight (1.03 kg; 95%CI 0.24; 1.82), waist to hip ratio 0.006 (95%CI 0.003; 0.011) and fasting glucose 0.09 mmol/L (95%CI 0.02; 0.15). The OR for COPD was 0.89 (95%CI 0.67; 1.18).

**Figure 4.**
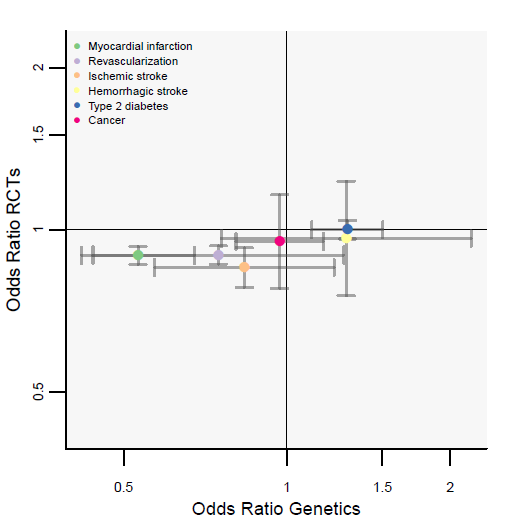
Associations of a *PCSK9* gene-centric score (GS) with non-cardiovascular events. Footnote: Effect estimates are presented as odds ratios (OR), with 95% confidence interval (CI) scaled to a 1 mmol/L lower LDL-C (mmol/L). Results are pooled using a fixed effect model. The size of the squares are proportional to the inverse of the variance. Note, that all GS estimates are based on 4 SNP, except for the Alzheimer’s disease estimate which excluded the SNP rs11591147 due to lack of data.

## Discussion

The genetic findings presented here show that variation in *PCSK9* is associated with lower circulating LDL-C and apoB concentrations, lower risk of MI and, with lesser confidence, the risk of ischemic stroke and coronary revascularization. These effects are consistent in direction to effects observed in the first PCSK9 inhibitor trial’s(17).

A recent systematic review of trial data(18) indicated a potential dysglycaemic effect from PCSK9 inhibition. PCSK9 monoclonal antibodies were associated with increased fasting glucose (0.17 as standardized mean difference [SMD] 95%CI 0.14; 0.19) and glycosylated haemoglobin (0.10 SMD 95%CI 0.07; 0.12)(18). Recently we, and others, showed natural genetic variation *PCSK9* was associated with elevated fasting glucose and T2DM(14, 19, 20) and that variation at other LDL-C-associated loci also influence risk of T2DM(21, 22). However, the FOURIER and ODYSSEY OUTCOMES trials, the largest treatment trials of PCSK9 inhibitors to date, did not find an association with risk of incident T2DM, at a median follow up of 2.2 and 2.8 years respectively. It is possible this reflects a genuine discordance between the findings from trials and genetic analyses. Alternatively, the exposure durations in the two largest trials may simply have been too short for subjects to develop T2DM. The risk increasing effect of statins on T2DM was only apparent after conducting a meta-analysis of 13 statin trials in which 4,278 T2DM cases were observed during an average follow-up of 4 years(23).

In general, inconsistencies between associations of variants in a gene encoding a drug target and the effects of the corresponding treatment are possible on a number of theoretical grounds. The effects of genetic variation (present from conception) may be mitigated by developmental adaptation or environmental changes. A null association of a genetic variant with an outcome therefore does not preclude an effect of a treatment administered in later life, when adaptive responses may no longer be available, or in the presence of a particular environment (24). We selected a subset of all genetic variants at *PCSK9* that capture information on many others and which have some annotated function. However, other approaches to more fully capture the entire gene-centric effect are worthy of future investigation(25).

The association of *PCSK9* variants with LDL-C and MI has been reported before(5), and was a motivating factor for the development of PCSK9 inhibiting drugs. Lotta and colleagues(19) reported a similar OR for MI of 0.60 (95%CI 0.48; 0.75) per 1 mmol/L decrease in LDL-C using the *PCSK9* rs11591147 SNP. Using a seven SNP *PCSK9* GS, Ference *et al*. reported a MI OR of 0.44 (95%CI 0.31; 0.64) per 1 mmol/L decrease in LDL-C(20). These scaled genetic effects are larger than the treatment effect observed in trials which others have noted previously(26), and ascribed to the lifelong effect of genetic variation versus the short-term effect of drug treatment in later life.

The available trial data showed PCSK9 inhibitors had a similar effect on MI (OR 0.90, 95%CI 0.86; 0.93) and ischemic stroke (OR 0.85 95%CI 0.78; 0.93). By contrast, the genetic analysis indicated a directionally concordant, but larger effect on MI (OR 0.53; 95%CI 0.42; 0.68) than ischemic stroke, (OR 0.84 95%CI 0.57; 1.22). The genetic analysis was, however, based on only 11,920 stroke cases, about one-fifth of the number of cases available for the genetic analysis of MI and as such confidence interval overlapped. We did observe a significantly differential association between *PCKS9* SNPs and ischemic and hemorrhagic stroke (interaction p-value = 0.02). Findings from statin trials previously suggested LDL-C lowering through inhibition of HMG-coA reductase is associated with a reduced risk of ischemic but potentially increased risk of hemorrhagic stroke(27–29). Our findings suggest that a different effect on ischemic and hemorrhagic stroke subtypes may be eventually identified for PCSK9 inhibitors.

Despite previous concerns about a potential effect of this class of drugs on cognition(30), the genetic analysis did not reveal a significant association of the four *PCSK9* variants with cognitive function or Alzheimer’s disease, nor with COPD or cancer, though this does not preclude an effect on such outcomes from drug treatment given in later life. While we explored the associations with any cancer (54702 events) as well as 12 individual cancer sites (Appendix Figure 2), we did not have data on some clinically relevant cancer types such as endometrial cancer.

This neutral effect on cognition has been recently reported by the EBBINGHAUS study, nested within the FOURIER trial, which reported a non-significant PCSK9 inhibitor effect on multiple measures of cognition confirming (using a non-inferiority design) an absence of effect(30); it should be noted that similar to the FOURIER and EBBINGHAUS (nested within the FOURIER) studies had similarly short follow-up. The absence of an effect on cognition during PCSK9 inhibitor treatment was also observed in the ODYSSEY OUTCOMES trial, which had a median follow-up(31) of 2.8 years.

Drugs (even apparently specific monoclonal antibodies) can exert actions on more than one protein if such targets belong to a family of structurally similar proteins. PCSK9, for example, is one of nine related proprotein convertases(32). Such ‘off-target’ actions, whether beneficial or deleterious, would not be shared by variants in the gene encoding the target of interest. In addition, monoclonal antibodies clear circulating PCSK9 from the blood and should not, in theory, influence any intracellular action of the protein(33).

Genetic association studies of the type conducted here tend to examine the risk of a first clinical event, whereas clinical trials such as ODYSSEY OUTCOMES focus on patients with established disease, where mechanisms may be modified. Proteins influencing the risk of a first event may also influence the risk of subsequent events, as observed in the case of the target of statin drugs that are effective in both primary and secondary prevention(1). For this and other reasons(34–36), examination of the effects of *PCSK9* variants on the risk of subsequent CHD events in patients with established coronary atherosclerosis is the subject of a separate analysis led by the GENIUS-CHD consortium(36).

In conclusion, *PCSK9* SNPs associated with lower LDL-C predict a substantial reduction in the risk of MI and concordant associations with a reduction in risk of ischemic stroke, but with a modestly increased risk of T2DM. We did not observe significant associations with other non-cardiovascular safety outcomes such as cancer, COPD, Alzheimer’s disease or atrial fibrillation. These findings suggest that it is likely that PCSK9 inhibitor therapy will not have any clinically meaningful effect on non-vascular diseases but it may increase risk of T2DM.

## Authors’ contributions

Amand F Schmidt, Daniel I Swerdlow, Michael V Holmes, Riyaz S Patel, Folkert W Asselbergs, Juan-Pablo Casas, Brendan J Keating, Aroon D Hingorani, David Preiss, Naveed Sattar contributed to the idea and design of the study. Amand F Schmidt, Daniel I Swerdlow, Michael V Holmes, designed the analysis scripts shared with individual centres. Amand F Schmidt performed the meta-analysis and had access to all the data. Amand F Schmidt, Aroon D Hingorani, and Juan-Pablo Casas drafted the initial manuscript. Michael V Holmes, Riyaz S Patel, Folkert W. Asselbergs, Brendan J Keating, David Preiss, Naveed Sattar, Daniel I Swerdlow, Zammy Fairhurst-Hunter, Fernando Pires Hartwig, Bernardo Lessa Horta, Elina Hypponen, Christine Power, Max Moldovan, Erik van Iperen, Kees Hovingh, Ilja Demuth, Kristina Norman, Elisabeth Steinhagen-Thiessen, Juri Demuth, Lars Bertram, Christina M Lill, Stefan Coassin, Johann Willeit, Stefan Kiechl, Karin Willeit, Dan Mason, John Wright, Richard Morris, Goya Wanamethee, Peter Whincup, Yoav Ben-Shlomo, Stela McLachlan, Jackie F. Price, Mika Kivimaki, Catherine Welch, Aida Sanchez, Pedro Marques-Vidal, Andrew Nicolaides, Andrie G. Panayiotou, N. Charlotte Onland-Moret, Yvonne T. van der Schouw, Giuseppe Matullo, Giovanni Fiorito, Simonetta Guarrera, Carlotta Sacerdote, Nicholas J Wareham, Claudia Langenberg, Robert A Scott, Jian’an Luan, Martin Bobak, Sofia Malyutina, Andrzej Pajak, Ruzena Kubinova, Abdonas Tamosiunas, Hynek Pikhart, Niels Grarup, Oluf Pedersen, Torben Hansen, Allan Linneberg, Tine Jess, Jackie Cooper, Steve E Humphries, Murray Brilliant, Terrie Kitchner, Hakon Hakonarson, David S. Carrell, Catherine A. McCarty, Kirchner, H Lester, Eric B. Larson, David R. Crosslin, Mariza de Andrade, Dan M Roden, Joshua C Denny, Cara Carty, Stephen Hancock, John Attia, Elizabeth Holliday, Rodney Scott, Peter Schofield, Martin O’Donnell, Salim Yusuf, Michael Chong, Guillaume Pare, Pim van der Harst, M. Abdullah Said, Ruben N. Eppinga, Niek Verweij, Harold Snieder, Tim Christen, D.O. Mook-Kanamori, Stefan Gustafsson, Lars Lind, Erik Ingelsson, Raha Pazoki, Oscar Franco, Albert Hofman, Andre Uitterlinden, Abbas Dehghan, Alexander Teumer, Sebastian Baumeister, Marcus Dörr, Markus M. Lerch, Uwe Völker, Henry Völzke, Joey Ward, Jill P Pell, Tom Meade, Ingrid E. Christophersen, Anke H. Maitland-van der Zee, Ekaterina V. Baranova, Robin Young, Ian Ford, Archie Campbell, Sandosh Padmanabhan, Michiel L Bots, Diederick E. Grobbee, Philippe Froguel, Dorothée Thuillier, Ronan Roussel, Amelie Bonnefond, Bertrand Cariou, Melissa Smart, Yanchun Bao, Meena Kumari, Anubha Mahajan, Paul M Ridker, Daniel I. Chasman, Alex P. Reiner, Marylyn D Ritchie, Leslie A Lange, Chris Finan, Ghazaleh Fatemifar, Rupert Faraway, Spiros Denaxas, Harry Hemingway, Richard Houlston, Alex J. Cornish, Sara E. Dobbins, Kari Hemminki, Ben Kinnersley, Marc Sanson, Karim Labreche, Matthias Simon, Melissa Bondy, Philip Law, Helen Speedy, James Allan, Ni Li, Molly Went, Niels Weinhold, Gareth Morgan, Pieter Sonneveld, Björn Nilsson, Hartmut Goldschmidt, Kari Hemminki, Amit Sud, Andreas Engert, Markus Hansson, Dennis Valentine, Jemma C. Hopewell, Sudha Seshadri, Caroline Dale, and Rui Providencia E Costa were responsible for study specific analyses and/or critically revised the manuscript.

## Guarantor

Amand F Schmidt performed the presented analyses, had full access to all the data in the study and takes responsibility for the integrity of the data and the accuracy of the data analysis.

